# Abstinence from cocaine self-administration promotes microglia pruning of astrocytes which drives cocaine-seeking behavior

**DOI:** 10.1101/2024.09.20.614128

**Authors:** Anze Testen, Jonathan W VanRyzin, Tania J Bellinger, Ronald Kim, Han Wang, Matthew J Gastinger, Emily A Witt, Janay P Franklin, Haley A Vecchiarelli, Katherine Picard, Marie-Ève Tremblay, Kathryn J Reissner

## Abstract

Rodent drug self-administration leads to compromised ability of astrocytes to maintain glutamate homeostasis within the brain’s reward circuitry, as well as reductions in surface area, volume, and synaptic colocalization of astrocyte membranes. However, the mechanisms driving astrocyte responses to cocaine are unknown. Here, we report that long-access cocaine self-administration followed by prolonged home cage abstinence results in decreased branching complexity of nucleus accumbens astrocytes, characterized by the loss of peripheral processes. Using a combination of confocal fluorescence microcopy and immuno-gold electron microscopy, we show that alterations in astrocyte structural features are driven by microglia phagocytosis, as labeled astrocyte membranes are found within microglia phagolysosomes. Inhibition of complement C3-mediated phagocytosis using the neutrophil inhibitory peptide (NIF) rescued astrocyte structure and decreased cocaine seeking behavior following cocaine self-administration and abstinence. Collectively, these results provide evidence for microglia pruning of accumbens astrocytes across cocaine abstinence which mediates cocaine craving.

## INTRODUCTION

Astrocytes are highly specialized glial cells characterized by their unique morphology, defined by thousands of fine, sheet-like processes that form distinct, bushy territories with minimal overlap between adjacent cells ^1–3^. These processes, which account for up to 95% of an astrocyte’s volume, contact numerous synapses and blood vessels, allowing astrocytes to adapt dynamically to changes in their environment ^4,5^. This intricate structural framework facilitates the ability of astrocytes to dynamically remodel their processes in response to fearful and sensorimotor stimuli, injury or inflammation, and in processes ranging from feeding, sleep, lactation, and cognition ^6–10^, effectively mediating a broad range of functions in health and disease. Among the stimuli that evoke structural responses of astrocytes, exposure to drugs of abuse leads to long-lasting changes in astrocyte structure and function ^11,12^. In particular, rat self-administration of cocaine leads to enduring deficits in regulation of glutamate homeostasis in the reward circuitry, which are also observed in human populations ^13,14^. Glutamate homeostasis is largely mediated by astrocytes, thereby directly implicating astrocytes in the pathophysiology of substance use disorders. Moreover, rat self-administration of drugs including cocaine, methamphetamine, and heroin, followed by a period of extinction or abstinence, is associated with an atrophic astrocyte phenotype, characterized by decreased astrocyte surface area, volume, and synaptic colocalization within the nucleus accumbens (NAc), a key region for reward processing ^15–19^. However, the mechanism of astrocyte atrophy and its relationship to the complexity of astrocyte branching remains unknown.

In the current study, we first sought to generate a deeper understanding of the nature of structural deficits observed in NAc astrocytes following cocaine self-administration and abstinence through investigation of process density, branching pattern and length. To this end, we developed and employed a three-dimensional analysis pipeline to reconstruct NAc astrocytes in two commonly used rodent models of drug abuse: short access self-administration (ShA) and extinction, and long-access self-administration (LgA) and prolonged abstinence. While reductions in astrocyte structural complexity were observed in both behavioral paradigms, these effects were significantly greater following long-access and prolonged abstinence. Interestingly, while overall astrocyte process density, branching pattern and number was reduced by cocaine abstinence, we found no effect on the average branch length, suggesting the loss of entire astrocyte processes ^20,21^.

Missing structural elements of astrocytes raised the possibility of pruning as a putative mechanism driving decreases in size and complexity, especially since cocaine has also been shown to induce structural changes in microglia, the brain’s resident innate immune cell. For example, two weeks of cocaine self-administration increases Iba1 staining and decreases microglial volume in the NAc as early as one day of withdrawal ^22,23^. Furthermore, cocaine can induce changes in microglia function via activation of microglia-expressed Toll Like Receptor 2 (TLR2) and 4 (TLR4) ^24^. Yet despite these reports that cocaine induces alterations in the glial landscape of the brain, very little is known about how these changes induce pathological plasticity and behavior that mediate relapse vulnerability.

Crosstalk and interactions between astrocytes and microglia were reported previously in multiple physiological and pathological conditions but mostly relied on cytokine and complement signaling pathway with no evidence of direct physical interaction ^25,26^. To this end, we sought to investigate evidence of microglia pruning of NAc astrocytes across cocaine abstinence. We found that long-access self-administration with prolonged abstinence increased microglia engulfment of astrocyte membranes. Moreover, inhibiting complement-dependent phagocytosis across abstinence rescued the cocaine-dependent reductions in astrocyte morphology, and decreased cocaine-seeking behavior. While phagocytic microglia are well described under both pathological conditions as well as development ^27^, this is to our knowledge the first observation of evidence for engulfment of astrocyte membranes by microglia. These observations further indicate that microglia phagocytosis is a contributing mechanism to protracted cocaine craving across abstinence.

## RESULTS

### Decreased Structural Complexity of NAc Astrocytes from Cocaine-Abstinent Rats

Abstinence from cocaine self-administration has been previously shown to induce an atrophic phenotype in NAc core astrocytes ^15,18,19^. To further explore whether this atrophy is associated with changes in astrocyte structural complexity, we conducted 3-dimensional reconstructions and filament tracing of Lck-GFP positive astrocytes across two behavioral paradigms (Figure 1A). Gross morphometric measurements of reconstructed astrocytes used for complexity analysis were published previously ^18,19^. These reconstructions were used as an input for Sholl analysis, which revealed significant differences in the number and distribution of Sholl intersections—used as a proxy for structural complexity—relative to the distance from the nucleus across the compared groups (Figure 1C).

**Figure 1.**
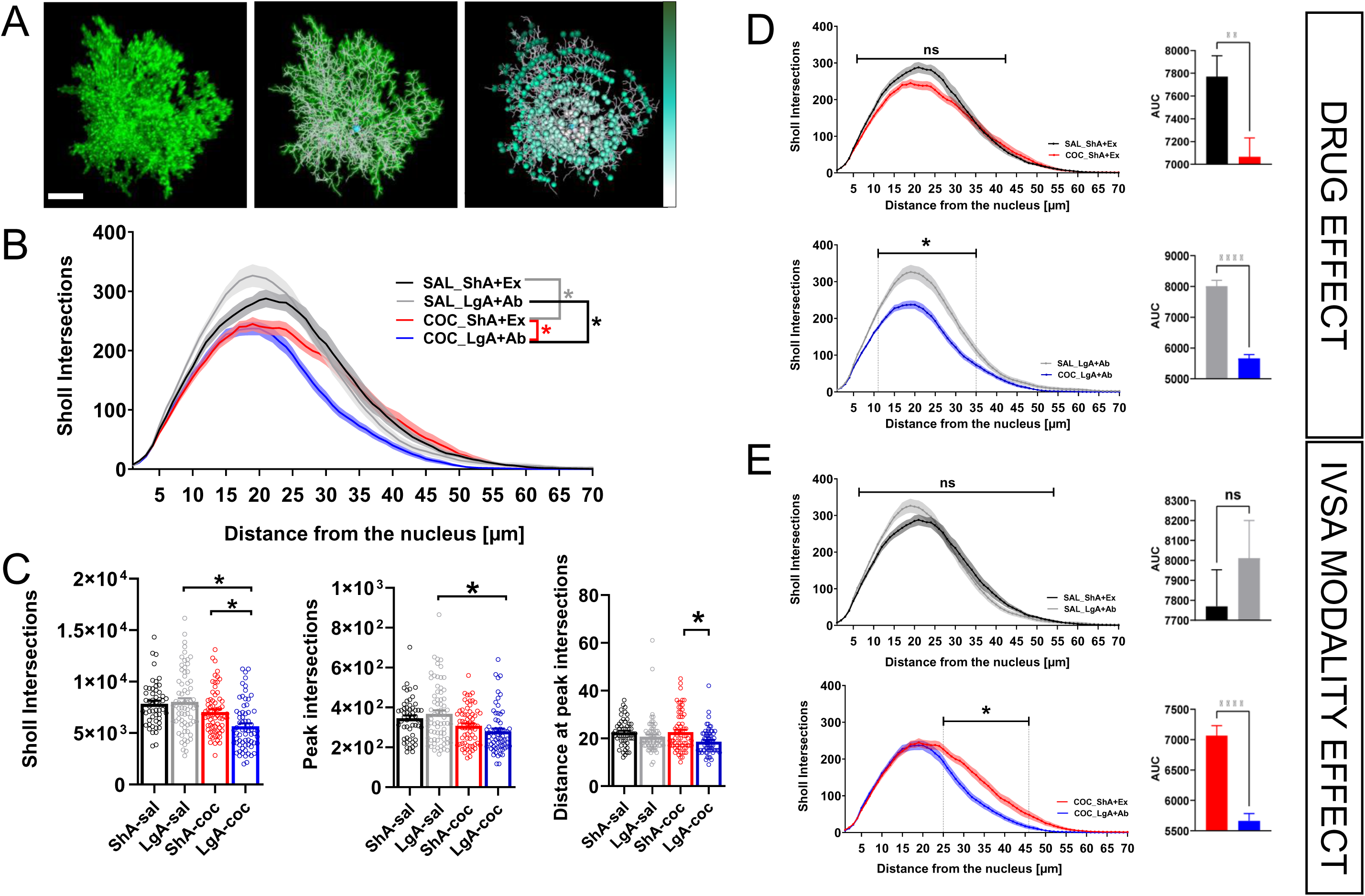
Cocaine induced atrophic phenotype of accumbal astrocytes is characterized by decreased structural complexity. (A) Timelines for each self-administration paradigm (B) 3-dimensional reconstructions of isolated masks of Lck-GFP positive astrocytes (left) were used for filament tracing to construct wire models of each astrocyte (middle) to be used for Sholl analysis (right). Scale bar: 20µm (C) Sholl intersections from all 4 groups plotted as a function of distance from the nucleus. Three-way interaction (distance x paradigm x drug: F(93, 2604) = 1.334, p=0.0194); two-way interactions (distance x paradigm: F(93, 2604) = 10.960, p<0.0001, distance x drug: F(93, 2604) = 9.383, p<0.0001). Three-way RM ANOVA. Asterixis signifies statistically significant group difference confirmed with 2-way RM ANOVA (expanded on in E and F). (D) The total number of Sholl intersections (left; F(3, 28)=9.111, p=0.0002), the peak intersections (maximum intersection value) (middle; F(3, 28)=5.342, p=0.0049)) and the distance at the peak complexity (right; F(3, 28)=4.908, p=0.0025) are all decreased in LgA-coc group. Each data point represents an analyzed cell (nested factor). One-way nested ANOVA with Tukey post-hoc test. (E) Sholl intersections, plotted as a function of distance for both behavioral paradigms separately and their respective AUC. Larger decrease in astrocyte complexity observed for the cocaine group with the LgA paradigm (bottom; main effect of the drug: F(1, 14)=20.62, p=0.0005; AUC: p<0.0001) then with the ShA group (top; main effect of the drug: F(1, 14)=1.802, p=0.2009; AUC: P=0.0079). 2-way RM ANOVA with Bonferroni post-hoc test (distance plots) and two-tailed unpaired t-test (AUC). (F) Sholl intersections, plotted as a function of distance for both drug conditions separately and their respective AUC. No effect of the self-administration paradigm on the saline control groups (top; main effect of the paradigm: F(1, 13)=0.1677, p=0.6888; AUC: p=0.03682) while LgA-coc shows more decrease in complexity compared to ShA-coc (bottom; main effect of the paradigm: F(1, 15)=9.734, p=0.0070; AUC: p<0.001). Two-way RM ANOVA with Bonferroni post-hoc test (distance plots) and two-tailed unpaired t-test (AUC). ns=not significant. Data are represented as mean +/- SEM. Significance was set at p<0.05. Group sizes: ShA-sal: N=7, n=6-9; ShA-coc: N=9, n=8; LgA-sal: N=8, n=8-11; LgA-coc: N=8, n=8-10 (N=number of animals, n=number of cells per animal). Detailed statistics for all the interactions in three- and two-way ANOVA reported in supp. Stats. Table #1. All group comparisons for the one-way nested ANOVA reported in the supp. Stats. Table #2.

We next more closely examined the contributions of cocaine across the two different self-administration models (Figure 1E) and assessed the influence of the behavioral paradigm under the same drug condition (Figure 1F). In the ShA paradigm, the cocaine-associated reduction in astrocyte structural complexity, when measured by distance from the nucleus, was not significant (Figure 1E, top). However, a significant decrease was observed in both the area under the curve (AUC) and the total number of intersections (Figure 1D, left). In contrast, the LgA paradigm revealed a marked reduction in astrocyte complexity across much of the cell’s length in the cocaine group, with a significant decrease between 11-35 µm from the nucleus (main effect of the drug: p=0.0005) (Figure 1E, bottom). This reduction was further reflected in decreased total intersections, peak intersections, and AUC (Figure 1D, left, middle), indicating a cell-wide atrophic phenotype characterized by reduced structural complexity specifically following the LgA/prolonged abstinence training.

To investigate the specific effect of the behavioral paradigm within the same drug condition, we compared saline control groups and found no significant differences in the distribution of Sholl intersections, total intersections, or peak intersections (Figure 1F, top; Figure 1D, left, middle), suggesting that the observed atrophy is primarily induced by cocaine. Furthermore, the LgA-cocaine group exhibited a more pronounced decrease in structural complexity than the ShA-cocaine group (Figure 1D, left). Notably, the reduction in total intersections in the LgA-cocaine group was mainly attributed to decreased complexity in distal astrocyte processes, with a significant reduction between 25-46 µm from the nucleus (p=0.0070) (Figure 1F, bottom). Additionally, the LgA-cocaine group exhibited an earlier peak in process complexity (density) compared to the ShA-cocaine group (Figure 1D, right), potentially indicating a shift in astrocyte process distribution following prolonged cocaine exposure.

Together, these results link the previously observed atrophic phenotype to a significant loss of structural complexity in astrocyte processes following cocaine self-administration and withdrawal, which is exacerbated in the LgA/prolonged abstinence cocaine group.

### Cocaine-Induced Decrease in the Structural Complexity of Accumbal Astrocytes Is Characterized by Attenuated Branching of Astrocyte Processes

The structural complexity of astrocytes, like other cell types, is largely governed by their branching patterns ^3^. Given the significant reduction in astrocyte structural complexity observed following cocaine self-administration, we performed a branching analysis to determine whether this reduction was due to a loss of specific branching patterns. We categorized bifurcation nodes into three subtypes: arborizing (bifurcation gives rise to two nodes that also branch), continuing (one node branches while the other terminates), and terminating (bifurcation gives rise to two terminating branches) (Figure 2A).

**Figure 2.**
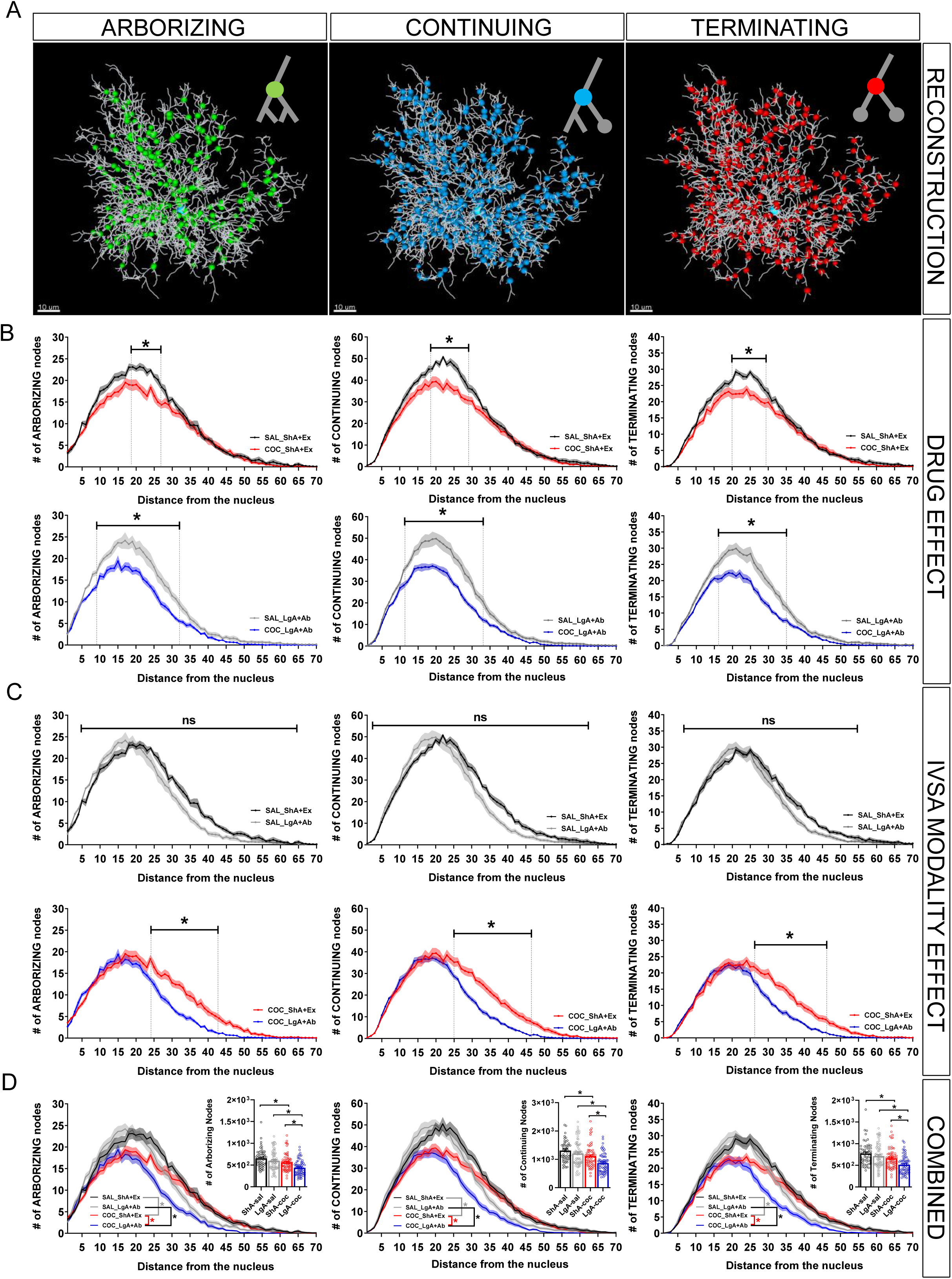
Cocaine-induced decrease in structural complexity of accumbal astrocytes results from decreased branching of peripheral processes. (A) Wire model of astrocyte processes showing arborizing (left), continuing (middle) and terminating (right) bifurcation nodes. Scale bar: 10µm (B) Number of bifurcations, plotted as a function of distance for both behavioral paradigms separately. ShA (top) (arborizing (left) - main effect of the drug: F(1, 14)=5.391, p=0.0358; continuing (middle) - main effect of the drug: F(1, 14)=6.072, p=0.0273; terminating (right) - main effect of the drug: F(1, 14)=7.529, p=0.0158). LgA (bottom) (arborizing (left) - main effect of the drug: F(1, 14)=19.20, p=0.0006; continuing (middle) - main effect of the drug: F(1, 14)=22.02, p=0.0003; terminating (right) - main effect of the drug: F(1, 14)=18.48, P=0.0007). Two-way RM ANOVA with Bonferroni post-hoc test. (C) Number of bifurcations, plotted as a function of distance for drug conditions separately. Saline (top) (arborizing (left) - main effect of the paradigm: F(1, 13)=2.343, p=0.1498; continuing (middle) - main effect of the paradigm: F(1, 13)=2.326, p=0.1512; terminating (right) - main effect of the paradigm: F(1, 13)=3.794, p=0.0734). Cocaine (bottom) (arborizing (left) - main effect of the paradigm: F(1, 15)=10.87, p=0.0049; continuing (middle) - main effect of the paradigm: F(1, 15)=11.31, p=0.0043; terminating (right) - main effect of the paradigm: F(1, 15)=10.71, p=0.0051). Two-way RM ANOVA with Bonferroni post-hoc test. ns=not significant. (D) Type-specific bifurcations of all 4 groups plotted together as a function of distance. Arborizing (left) – distance x paradigm: F(91, 2548)=9.709, p<0.0001; distance x drug: F(91, 2548)=8.143, p<0.0001. Continuing (middle) - distance x paradigm: F(91, 2548)=9.772, p<0.0001; distance x drug: F(91, 2548)=10.330, p<0.0001. Terminating (right) - distance x paradigm: F(91, 2548)=6.510, p<0.0001; distance x drug: F(91, 2548)=9.617, p<0.0001. Three-way RM ANOVA. Asterixis signifies statistically significant group difference confirmed with two-way RM ANOVA. Panel inserts (total number of type-specific bifurcations): arborizing (left) – F(3, 28)=9.607, p=0.0002; continuing (middle) – F(3, 28)=10.10, p=0.0001; terminating (right) – F(3, 28)=9.093, p=0.0002. One-way nested ANOVA with Tukey post-hoc test. Data are represented as mean +/- SEM. Significance was set at p<0.05. Group sizes: ShA-sal: N=7, n=6-9; ShA-coc: N=9, n=8; LgA-sal: N=8, n=8-11; LgA-coc: N=8, n=8-10 (N=number of animals, n=number of cells per animal). Detailed statistics for all the interactions in three- and two-way ANOVA reported in supp. Stats. Table #1. All group comparisons for the one-way nested ANOVA reported in the supp. Stats. Table #2.

In astrocytes following the ShA/extinction paradigm, we observed a significant decrease in branching at the peak of branching density across all three bifurcation subtypes: arborizing (18-27 µm; main effect of the drug: p=0.0358), continuing (18-28 µm; p=0.0273), and terminating (20-29 µm; p=0.0158) nodes compared to saline controls (Figure 2B, top). In contrast, the LgA/abstinence cocaine group showed impaired branching across a broader region of the cell: arborizing (8-32 µm; p=0.0006), continuing (11-33 µm; p=0.0003), and terminating (16-35 µm; p=0.0007) nodes (Figure 2B, bottom).

Next, we evaluated the impact of the behavioral paradigm by comparing the saline control groups and found no significant changes in branching profiles, confirming that cocaine is necessary for inducing the atrophic phenotype and impairing astrocyte structural complexity (Figure 2C, top). When comparing the cocaine-treated groups, the LgA group exhibited a more pronounced reduction in distal branching compared to the ShA group: arborizing (24-43 µm; p=0.0049), continuing (25-46 µm; p=0.0043), and terminating (26-29 µm; p=0.0051) nodes. These results suggest that distal astrocyte processes are more vulnerable to atrophy after prolonged cocaine exposure and withdrawal.

The overall reduction in astrocyte structural complexity was also reflected in the total number of bifurcations within each paradigm and drug condition (Figure 2D – insets), as well as in the combined total bifurcations across subtypes (Figure S1A). Notably, all bifurcation subtypes showed similar decreases, indicating that the mechanism of reduction is not subtype-specific. Interestingly, continuing nodes accounted for approximately 50% of all bifurcations in each group (ShA-sal: 47.90%, ShA-coc: 47.73%, LgA-sal: 47.85%, LgA-coc: 47.66%), and this ratio remained consistent despite the cocaine-associated reductions (Figure S2B), suggesting a global mechanism of structural impairment rather than one confined to a specific bifurcation type.

### Loss of Peripheral Astrocyte Processes (PAPs) as a Hallmark of the Cocaine-Associated Atrophic Phenotype

The structural deficits observed in astrocytes, including reductions in process density and branching, can be largely attributed to the loss of peripheral astrocyte processes (PAPs), particularly in the distal regions of the cells. PAPs are critical functional units of astrocytes ^3^ and are known for their structural plasticity in both health and disease ^28,29^. To determine whether the PAPs were retracted or entirely lost following the different behavioral paradigms, we used terminal points of our reconstructed filaments as proxies for PAPs, since they are often not resolvable using conventional light microscopy ^30^ (Figure 3A).

**Figure 3.**
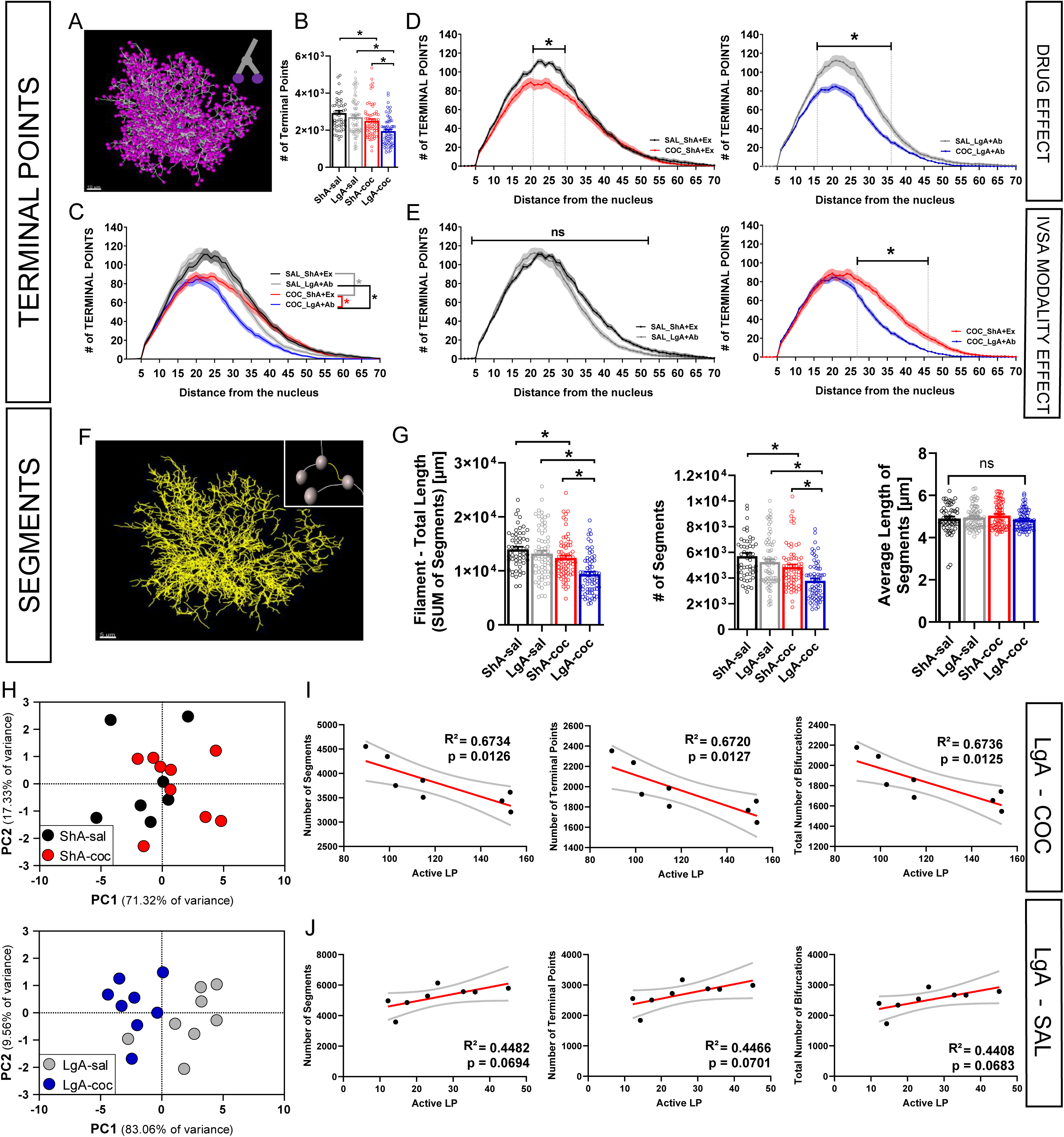
The cocaine-induced decrease in branching of accumbal astrocytes is a result of a loss of peripheral astrocyte processes. (A) Wire model of reconstructed astrocyte showing terminal points of most peripheral segments (a proxy for PAPs). Scale bar: 10µm (B) Total number PAPs for all groups (F(3, 28)=9.297, p=0.0002). One-way nested ANOVA with Tukey post-hoc test. (C) Number of PAPs for all 4 groups plotted as a function of distance from the nucleus (distance x paradigm: F(91, 2548)=8.012, p<0.0001; distance x drug: F(91, 2548)=9.953, p<0.0001). 3-way RM ANOVA. Asterixis signifies statistically significant group difference confirmed with 2-way RM ANOVA. (D) Number of PAPs, plotted as a function of distance for both behavioral paradigms separately (ShA (left) - main effect of the drug: F(1, 14)=5.287, p=0.0374; LgA (right) - main effect of the drug: F(1, 14)=19.71, p=0.0006). Two-way RM ANOVA with Bonferroni post-hoc test. (E) Number of PAPs, plotted as a function of distance for both drug conditions separately (saline (left) - main effect of the paradigm: F(1, 13)=1.842, p=0.1978; cocaine (right) - main effect of the paradigm: F(1, 15)=10.80, p=0.0050). 2-way RM ANOVA with Bonferroni post-hoc test. ns = not significant (F) A complete wire model of an isolated astrocyte, used to measure the combined length of the filament and count its segments (Insert: a single segment; defined as a part of the filament between two bifurcation nodes). Scale bar: 5µm (G) Total length of the filament (F(3, 28)=12.19, p<0.0001), number of segments (F(3, 28)=9.626, p=0.0002) and average length of the segments (F(3, 28)=0.4517, p=0.7181) for all groups. One-way nested ANOVA with Tukey post-hoc test. ns = not significant (H) Principle component analysis of all the measured astrocyte complexity parameters shows clear separation and clustering between the cocaine and saline groups for LgA (bottom), with the PC1 contributing to the separation of the groups (PC1: POV=83.06%, Eigenvalue=9.137; PC2: POV=9.56%, Eigenvalue=1.052; PCR (ALP): F(2, 13)=10.14, P=0.0022) while in the ShA paradigm (top), the separation is not clear (PC1: POV=71.32%, Eigenvalue=7.846; PC2: POV=17.33%, Eigenvalue=1.906; PCR (ALP): F(2, 13)=2.837, p=0.0949). PCs used in PCR selected by the Keiser rule. (I) Regression analysis for the LgA-coc group, showing correlation between the number of active lever presses (ALP) during SA and most relevant complexity parameters of astrocytes: number of segments (left - R2=0.6734, F(1, 6)=12.37, p=0.0126), number of PAPs (middle - R2=0.6720, F(1, 6)=12.29, p=0.0127) and the total number of all bifurcations nodes (right - R2=0.6736, F(1, 6)=12.38, p=0.0125). Plotted as a simple linear regression with the 95% confidence interval. (J) Simple liner regression analysis for the LgA-sal group, showing the lack of correlation between active lever presses (ALP) during SA and most relevant complexity parameters of astrocytes: number of segments (R2=0.4482, F(1, 6)=4.841, p=0.0694), number of PAPs (R2=0.4466, F(1, 6)=4.841, p=0.0701) and the total number of all bifurcations nodes (R2=0.4508, F(1, 6)=4.925, p=0.0683). Plotted as a simple linear regression with the 95% confidence interval. Plotted as a simple linear regression with the 95% confidence interval. Data are represented as mean +/- SEM. Significance was set at p<0.05. Group sizes: ShA-sal: N=7, n=6-9; ShA-coc: N=9, n=8; LgA-sal: N=8, n=8-11; LgA-coc: N=8, n=8-10 (N=number of animals, n=number of cells per animal). Detailed statistics for all the interactions in three- and two-way ANOVA reported in supp. Stats. Table #1. All group comparisons for the one-way nested ANOVA reported in the supp. Stats. Table #2. See SF2 for PCA analysis loadings, biplots and full POV.

Similar to our findings with Sholl intersections and bifurcations, cocaine self-administration led to a reduction in the total number of terminal points (Figure 3B). In the ShA group, there was a significant decrease in terminal points at the peak density (21-29 µm; p=0.0374) in cocaine versus saline, while the LgA cocaine group showed deficits across a broader portion of the cell compared to LgA saline (16-36 µm; p=0.0006) (Figure 3D). Furthermore, the LgA group exhibited a more substantial loss of distal PAP proxies compared to the ShA group (27-46 µm; p=0.0050) (Figure 3E). No significant differences in terminal point distribution were observed between the two saline control groups (Figure 3E, left).

To confirm that these reductions in astrocyte process density and number were due to process loss rather than shrinkage, we analyzed the full filament reconstructions and quantified the lengths of branch segments between bifurcating nodes (Figure 3F, inset). Consistent with previous results, we observed a reduction in both the total number and the total length of segments in the cocaine groups compared to their saline controls, with the LgA group showing a more pronounced decrease than the ShA group (Figure 3G, left, middle). Surprisingly, the average length of the segments remained consistent across all behavioral paradigms and drug conditions (Figure 3G, right), definitively indicating that the atrophic phenotype caused by cocaine is due to astrocyte process loss rather than shrinkage.

Next, we investigated whether astrocyte complexity profiles could distinguish between groups based on their self-administration history. A principal component analysis (PCA) revealed that two principal components explained 71.32% and 17.33% of the variance for ShA animals (saline and cocaine) and 83.06% and 9.56% for LgA animals (saline and cocaine). Interestingly, the ShA group did not show obvious clustering, suggesting that deficits in astrocyte complexity in this group were not severe enough to predict group identity (Figure 3H, top). In contrast, the LgA group showed clear clustering, indicating that complexity measurements are strong predictors of group identity and could potentially serve as biomarkers for the cocaine-induced atrophic phenotype (Figure 3H, bottom).

Finally, we explored whether astrocyte complexity measurements correlated with cocaine-seeking behavior. Focusing on the LgA paradigm, which showed predictive value in the PCA, we performed a linear correlation analysis using parameters that had shown significant correlations in a parallel PCR (principal component regression) analysis. All selected complexity parameters (number of terminal points, bifurcations, and segments) were significantly correlated with the number of active lever presses during cocaine self-administration, directly linking the magnitude of the atrophic phenotype to cocaine-seeking behavior (Figure 3I). As expected, the LgA-saline group showed no correlation between structural complexity and seeking behavior (Figure 3J).

In summary, these findings suggest that the loss of peripheral astrocyte processes is the primary factor responsible for the decrease in astrocyte structural complexity observed following cocaine self-administration and withdrawal. This atrophic phenotype distinguishes cocaine-exposed astrocytes from those in a healthy brain.

### Microglial Morphology Is Not Affected by Cocaine Self-Administration and Withdrawal

To determine whether astrocytes are the only glial subtype affected by cocaine self-administration (SA) and withdrawal, we next examined microglia, which have also been implicated in habitual drug-taking behavior ^31^. We focused entirely on cellular features following LgA self-administration, as it produced the most robust atrophic phenotype in astrocytes, characterized by decreased structural complexity (LgA = WD45). Microglia features were assessed by antibody labeling for Iba1 and confocal microscopy on either withdrawal day 1 (WD1) or 45 following cocaine or saline self-administration (Figure 4A).

**Figure 4.**
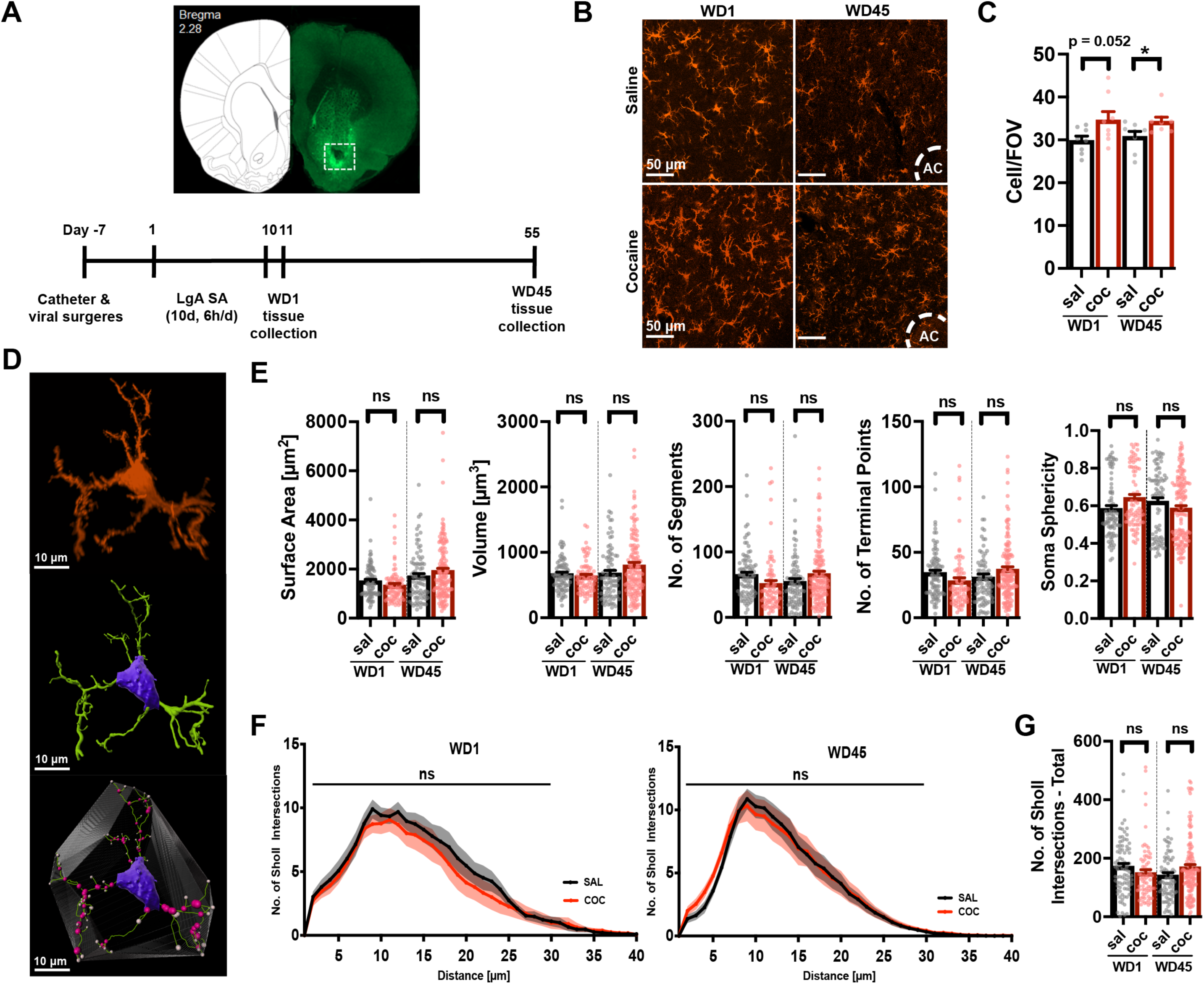
There were no detected microglia morphological changes at WD1 or WD45 following LgA cocaine self-administration. There was a significant increase in the number of microglia in the nucleus accumbens at both time points. (A) Whole-brain slice showing the nucleus accumbens, with expression of AAV5-GfaABC1D-Lck-GFP (top). Microglia were examined from within the white square. Timeline of self-administration and tissue collection points for microglia analysis (bottom). (B) Microglial cells were counted at WD1 and WD45. The images on the right depict 20x representative fields of view (FOV) images of Iba1-stained microglia. (C) Cell counts were measured at each time point by looking at cell/FOV. WD1 sal vs. coc F(1, 14)=2.115, p=0.0528; WD45 sal vs. coc F(1, 14)=2.272, p= 0.0394. (D) Individual microglial cell stained with Iba1 (top). 3-dimensional reconstruction of the same cell using Bitplane Imaris Software showing the soma in purple and reconstruction of processes in green (middle). The reconstructed cell shows branch points (magenta) and terminal points (white) (bottom). (E) Microglial morphometric measurements were collected, and a few chosen graphs are shown from left to right: Surface area, volume, number of segments, number of terminal points, and soma sphericity. Surface area: sal vs. coc, WD1: F(1, 14)=0.8313, p=0.4197; WD45:F(1, 18)=0.06991, p=0.9450. Volume: sal vs. coc, WD1: F(1, 14)=0.5609, p=0.5837; WD45: F(1, 18)=0.1614, p=0.8736. Number of segments sal vs. coc, WD1: F(1, 14)=1.017, p=0.3265; WD45 F(1, 18)=0.5116, p=0.6152. Number of terminal points sal vs. coc, WD1: F(1, 14)=0.9016, p= 0.3825; WD45 F(1, 18)=0.4314, p= 0.6713. Soma sphericity: sal vs. coc, WD1: F(1, 14)=1.735, p= 0.2089; WD45 F(1, 18)=0.9626, p= 0.3485. (F) Sholl analysis was also conducted with shown graphs of WD1 and WD45. (G) The accompanying quantification for sholl analysis is shown in the graph depicted by total number of sholl intersections. * p < 0.05 between groups, all error bars are standard error of the mean (SEM). Data are represented as mean +/- SEM. Significance was set at P<0.05. Group sizes: WD1-sal: N=8, n=8-14; WD1-coc: N=8, n=7-17; WD45-sal: N=10, n=4-14; WD45-coc: N=10, n=4-26 (N=number of animals, n=number of cells per animal).

We first quantified the number of microglia in the NAc core and observed an increase in both the WD1 and WD45 cocaine-administering groups compared to saline controls (Figure 4B, 4C). Next, similar to our approach with astrocytes, we reconstructed filament models of individual microglia to assess potential changes in their structural complexity (Figure 4D). Interestingly, we found no significant changes in any morphometric parameters or complexity measurements between the cocaine-administering groups and their respective controls (Figure 4E, S4A). Likewise, no changes were observed in the number of microglial segments, branching points, or terminal points (Figure 4E, S4A), contrasting sharply with the astrocyte results. We also analyzed microglial somata (area, volume, and sphericity), which are thought to reflect the reactive state of microglia, but none of these measurements were significantly altered between groups (Figure 4E, S4B). To further investigate microglial structural complexity, we performed Sholl analysis on the reconstructed microglial models. Again, no significant changes were found in the distribution of Sholl intersections along the length of the cells or in the total number of Sholl intersections (Figure 4F,G). Finally, a principal component analysis (PCA) using all collected microglial structural data failed to produce predictive clustering (Figure S4C). Together, these results demonstrate that, unlike astrocytes, prolonged cocaine intake and abstinence do not induce morphological changes in microglia, suggesting that astrocytes are uniquely vulnerable to cocaine exposure.

### Peripheral Astrocyte Processes Are Associated with Phagocytic Microglia Following Prolonged Cocaine Self-Administration and Withdrawal

Even though we observed no significant morphological changes in microglia following cocaine self-administration (SA) and withdrawal, we did detect an increased number of microglia in the NAc core at the WD45 time point. Microglia, as resident macrophages of the central nervous system, are primarily responsible for surveilling their environment and, upon activation, removing potential triggers ^32^. Activated microglia express various markers, including CD68, which is localized in microglial phagosomes, late endosomes, and lysosomes^33^.

To investigate whether cocaine exposure led to the activation of microglia, we performed colocalization analysis between CD68 and the microglial structural marker Iba1 (Figure 5A). CD68 expression was low in both saline controls and the ShA/extinction cocaine group, whereas it was markedly increased in the LgA/ abstinence cocaine group (Figure 5B, top). Due to the presence of virally expressed Lck-GFP specifically in astrocytes, we were able to observe an increased association between Lck-GFP-positive astrocyte membranes and Iba1-positive microglia in the LgA/abstinence cocaine group (Figure 5A). Colocalization analysis confirmed a significant increase in this association compared to saline controls, particularly at the WD45 time point (Figure 5B, middle). In summary, microglia in the NAc core following prolonged cocaine self-administration and withdrawal exhibit increased expression of the phagocytic marker CD68 and enhanced association with putative astrocyte processes, suggesting that microglia may be actively engaged in the clearance of astrocyte material.

**Figure 5.**
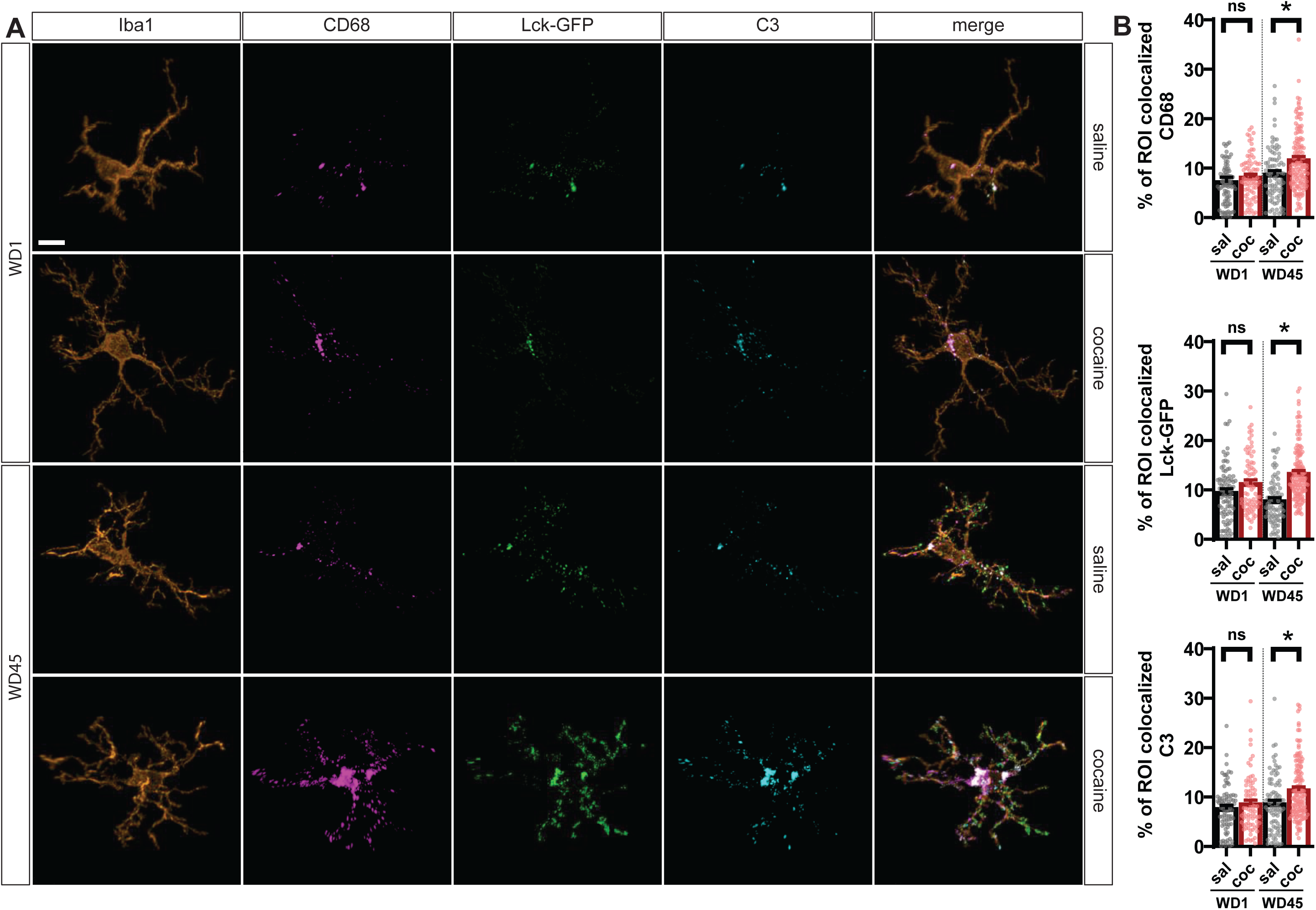
Cocaine SA and withdrawal leads to increased astrocytes-microglia interactions. (A) Representative images of isolated microglia (Iba1) from two SA paradigms, microglia-associated astrocyte processes (Lck-GFP) as well as microglia-colocalized CD68 and C3 signal. (B) Colocalization analysis for Iba1/CD68 (top – WD1: F(1, 14)=0.6292, p=0.4409; WD45: F(1, 18)=7.215, p=0.0151), Iba1/Lck-GFP (middle – WD1: F(1, 14)=1.950, p=0.1844; WD45: F(1, 18)=13.98, p=0.0015) and Iba1/C3 (bottom – WD1: F(1, 14)=0.9589, p=0.3441; WD45: F(1, 18)=5.242, p=0.0343). Two-tailed nested t-test. ns = not significant Data are represented as mean +/- SEM. Significance was set at p<0.05. Group sizes: WD1-sal: N=8, n=8-14; WD1-coc: N=8, n=7-17; WD45-sal: N=10, n=4-14; WD45-coc: N=10, n=4-26 (N=number of animals, n=number of cells per animal).

### Increase in Microglia-PAP Colocalization Following Cocaine Self-Administration and Prolonged Abstinence Is a Result of Microglial Phagocytosis of Peripheral Astrocyte Processes

To further investigate the unexpected association of astrocyte processes with phagocytic microglia, we generated orthogonal slices of reconstructed microglia. These revealed that Lck-GFP-positive material was located within the microglial soma (Figure 6A, left). This was corroborated by 3-dimensional reconstructions of Iba1-positive microglia (Figure 6A, right), which showed Lck-GFP-positive inclusions not only within the soma (red squares) but also within peripheral microglial processes (blue squares). To substantiate these observations, we employed transparent surface visualization for Iba1-positive microglia and CD68-positive phagosomes. This analysis confirmed that a subset of Lck-GFP-positive inclusions was located within CD68-positive phagosomes or late lysosomes (Figure 6C – I, II, III). These findings were further confirmed using scanning electron microscopy (SEM), which demonstrated Lck-GFP-positive inclusions within microglial vesicles (Figure 6C – IV, V).

**Figure 6.**
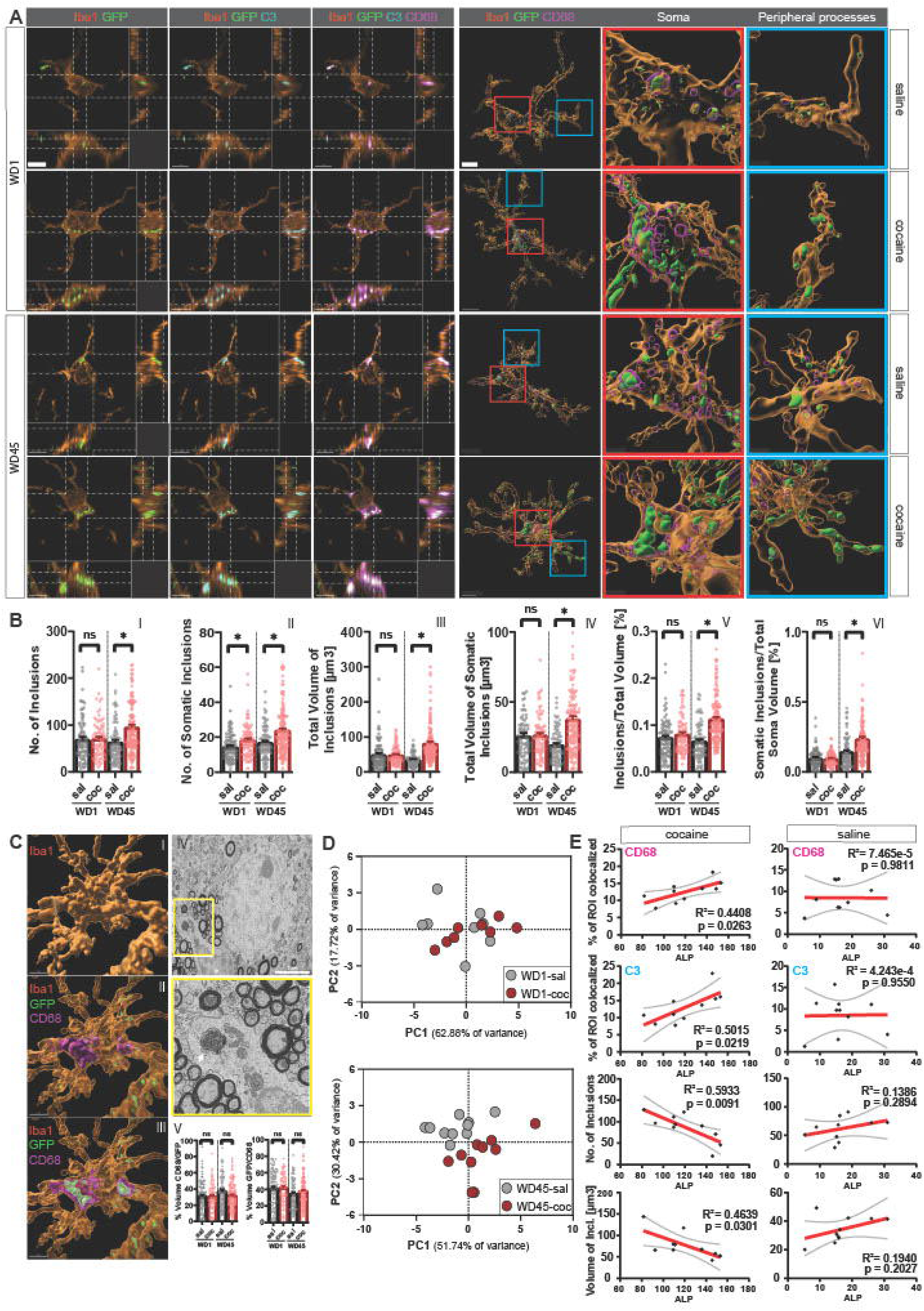
Increase in the microglia-PAPs colocalization following cocaine SA and prolonged abstinence is a result of microglial phagocytosis of PAPs FOV size. (A) Orthogonal projections of isolated microglia showing Iba1/Lck-GFP, Iba1/Lck-GFP/C3 and Iba1/Lck-GFP/C3/CD68 colocalization, respectively (right). The same isolated microglia, 3-D reconstructed with transparent surface, showing that colocalized GFP and CD68 signals are located inside the microglial cytoplasm and not merely colocalized with the cell surface/membrane. CD68- and GFP-positive microglial inclusions can be found in soma (red boxes) as well as in the peripheral microglial processes (blue boxes). (B) The number of total GFP-positive inclusions per microglia (I – WD1: F(1, 14)=0.0294, p=0.8663; WD45: F(1, 18)=6.076, p=0.0240), the number of GFP-positive inclusions per microglial soma ((II – WD1: F(1, 14)=5.970, p=0.0284; WD45: F(1, 18)=4.837, p=0.0412), total volume of GFP-positive inclusions per microglia (III – WD1: F(1, 14)=0.0133, p=0.9098; WD45: F(1, 18)=17.26, p=0.0006), total volume of GFP-positive inclusions per microglial soma (IV – WD1: F(1, 14)=0.1626, p=0.6928; WD45: F(1, 18)=14.61, p=0.0012), percentage of the total microglia volume occupied by the GFP-positive inclusions (V – WD1: F(1, 14)=1.067, p=0.3192; WD45: F(1, 18)=16.00, p=0.0008) and percentage of the total microglia soma volume occupied by the somatic GFP-positive inclusions (VI – WD1: F(1, 14)=0.0002, p=0.9890; WD45: F(1, 18)=35.01, p=0.001. Two-tailed nested t-test. ns = not significant (C) 3-dimensional reconstruction of a representative microglia (WD45-coc group) showing opaque surface of the soma and primary microglial processes (I). The same reconstruction with transparent surface, showing CD68-positive phagosome with opaque surface inside microglial soma (II). The same reconstruction with the transparent surface of the CD68-postive phagosome, showing GFP-positive astrocyte inclusions inside microglial phagosome (III). Scale bar. SBF-TEM micrography of a microglial phagosome showing GFP-positive inclusion (IV) Percentage of volume of all CD68-positive astrocyte inclusions versus total volume of GFP-positive astrocyte inclusions (left – WD1: F(1, 14)=0.7203, p=0.7203; WD45: F(1, 18)=1.389, p=0.2540) and a percentage of CD68-positive phagosome volume filed with Lck-GFP-positive inclusions versus total CD68-positive phagosome volume (right - WD1: F(1, 14)=0.2229, p=0.6441; WD45: F(1, 18)=0.5687, p=0.4605) (V). Two-tailed nested t-test. ns = not significant (D) Principle component analysis of all the measured microglial astrocyte-derived inclusion parameters shows clear separation and clustering between the cocaine and saline groups for the WD45 group (bottom), with both PCs contributing to the separation of the groups (PC1: POV=51.74%, Eigenvalue=5.691; PC2: POV=30.42%, Eigenvalue=3.346; PCR (infusions): F(2, 17)=17.47, p<0.0001) while at the WD1 timepoint (top), the separation is not clear (PC1: POV=62.88%, Eigenvalue=6.916; PC2: POV=17.72%, Eigenvalue=1.949; PCR not performed). PCs used in PCR selected by the Keiser rule. (E) Linear regression analysis for the WD45-cocaine group (left), showing correlation between the number of active lever presses (ALP) during SA and astrocyte inclusion-related parameters of microglia: CD68/Iba1 colocalization (R2=0.4804, F(1, 8)=7.397, p=0.0263), and C3/Iba1 colocalization (R2=0.5015, F(1, 8)=8.049, p=0.0219) positively correlated to ALP (top), while the total number of astrocyte-derived inclusions (R2=0.5933, F(1, 8)=11.670, p=0.0091) and the total volume of inclusions (R2=0.4639, F(1, 8)=6.922, p=0.0301) were correlated inversely (bottom). None of the inclusion measurements were correlated in the WD45-saline group (right): CD68/Iba1 colocalization (R2=7.465×10-5, F(1, 8)=0.0006, p=0.9811), C3/Iba1 colocalization (R2=0.0004, F(1, 8)=0.0033, p=0.9550, the total number of astrocyte-derived inclusions (R2=0.1386, F(1, 8)=1.287, p=0.2894), the total volume of inclusions (R2=0.1940, F(1, 8)=1.925, p=0.2027). Plotted as a simple linear regression with the 95% confidence interval. Data are represented as mean +/- SEM. Significance was set at p<0.05. Group sizes: WD1-sal: N=8, n=8-14; WD1-coc: N=8, n=7-17; WD45-sal: N=10, n=4-14; WD45-coc: N=10, n=4-26 (N=number of animals, n=number of cells per animal).

Quantification of Lck-GFP-positive inclusions within Iba1-positive microglia revealed a significant increase in the number of inclusions (I, II), their volume (III, IV), and the ratio of inclusion volume to total microglia volume (V, VI) in both the microglial soma and peripheral processes in the WD45-cocaine group compared to controls (Figure 6B). No significant changes were observed in the WD1 group. Additionally, we quantified the percentage of the total volume of all CD68-positive astrocyte inclusions relative to the total volume of GFP-positive astrocyte inclusions (CD68+GFP+ incl. / CD68-GFP+ incl.) and found no significant differences between groups (WD1-sal: 29.48%, WD1-coc: 31.34%, WD45-sal: 37.20%, WD45-coc: 31.95%). Similarly, the percentage of CD68-positive phagosome volume filled with Lck-GFP-positive inclusions relative to total CD68-positive phagosome volume (GFP+CD68+ incl. / GFP-CD68+ incl.) was consistent across groups (WD1-sal: 39.81%, WD1-coc: 42.70%, WD45-sal: 34.36%, WD45-coc: 37.87%) (Figure 6B – V). Despite the increase in Lck-GFP-positive inclusions in the WD45-cocaine group, these inclusions were equally distributed between CD68-positive phagosomes and cytoplasm/CD68-negative vesicles, suggesting that the increase in inclusions is not limited to a specific subcellular compartment.

Given the absence of any morphological differences in microglia, we sought to determine whether inclusion-related measurements could predict drug and control group identity. To test this, we performed a blinded PCA using all available inclusion-related measurements. The WD1 group did not show obvious clustering (Figure 6D, top), while the WD45 group exhibited tight clustering based on drug condition, with clear separation along both principal components (Figure 6D, bottom). This indicates that measurements of astrocyte-derived microglial inclusions are sufficient to distinguish between groups following prolonged abstinence from cocaine self-administration. Finally, we performed a linear regression analysis using ALP on inclusion parameters that were most significantly correlated based on a parallel PCR analysis with the PCA. CD68 exhibited a strong positive correlation with ALP, while the total number of inclusions (both in the soma and periphery) and the total volume of inclusions showed an inverse correlation with ALP. These findings suggest a novel glia-glia interaction, involving microglial phagocytosis of peripheral astrocyte processes following prolonged cocaine self-administration and withdrawal, which may contribute to the atrophic phenotype of astrocytes observed in this study.

### Blocking Microglia-Mediated Astrocyte Pruning Restores Astrocytic Synaptic Coverage and Decreases Cocaine Seeking

To test the hypothesis that microglia-mediated phagocytosis of astrocyte processes leads to structural deficits in astrocytes following cocaine self-administration and withdrawal, we bilaterally infused AAV5-GfaABC1D-Lck-GFP to label astrocytes and implanted guide cannulas targeting the NAc core. All animals underwent LgA cocaine self-administration, and beginning 24 hours after the last session, animals received microinjections of either the CR3 blocker neutrophil inhibitor factor (NIF) or sterile PBS as a control once weekly during home cage abstinence (Figure 7A,B). CR3 is expressed on microglia and serves as the receptor for the C3 ligand. By blocking the ability of microglia to recognize the C3 “eat me” signal, we predicted that NIF treatment would rescue the astrocyte morphological and deficits typically observed following abstinence from cocaine.

**Figure 7.**
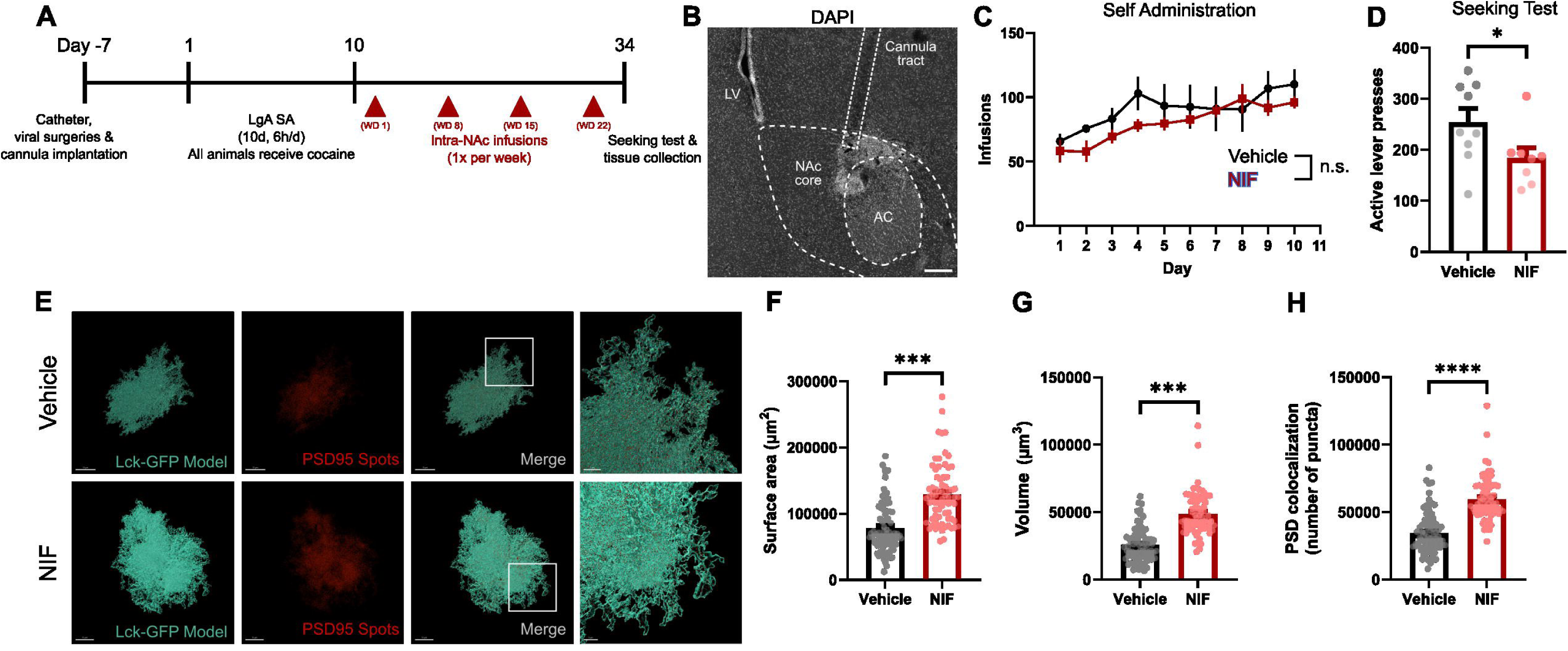
Blocking microglia-mediated astrocyte pruning restores astrocytic synaptic coverage and decreases cocaine seeking. (A) Timeline for self-administration and intra-NAc NIF infusions (B) Representative image of the NAc core showing location of cannula placement. Scale bar: 100 µm (C) The number of cocaine infusions across 10 days of LgA self-administration, showing no difference between groups (main effect of treatment: F(1, 15) = 1.069, p=0.3176) (D) The number of lever presses during cocaine seeking test on WD24. Animals that received intra-NAc infusions of NIF peptide once weekly during abstinence had a significant reduction in the number of lever presses (Mann Whitney U=14.50, p=0.038) (E) Representative 3D surface model of an astrocyte from vehicle (top left) or NIF (bottom left) treated animals with corresponding PSD95 spots model (middle left, top and bottom). Merged images show colocalization of PSD95 signal to astrocyte surface models (middle right, top and bottom). Inset indicates higher magnification image showing colocalization of PSD95 spots to perisynaptic astrocyte processes for vehicle and NIF-treated animals (right, top and bottom). Scale bars: 15 µm and 5 µm (inset) (F) Astrocyte surface area quantified in vehicle and NIF-treated animals. Nested t test t(17) = 4.343, p < 0.001 (G) Astrocyte volume quantified in vehicle and NIF-treated animals. Nested t test t(17) = 5.002, p < 0.001 (H) Number of PSD95 spots colocalized to astrocytes in vehicle and NIF-treated animals. Nested t test t(17) = 5.360, p < 0.001 Data are represented as mean +/-SEM. Significance was set at p<0.05. Group sizes: vehicle: N=9-10, n=7-9; NIF: N=8-9, n=8; (N=number of animals, n=number of cells per animal).

As expected, all animals underwent cocaine self-administration with no significant differences in drug-taking behavior prior to the intervention (Figure 7C). Forty-eight hours following the final NIF administration, the animals underwent a cocaine-seeking test. The NIF-treated group showed a significant reduction in cocaine-seeking behavior compared to PBS-treated controls (Figure 7D).

To confirm that the reduction in cocaine seeking was related to the rescue of astrocyte morphology, we collected tissue samples immediately following the seeking test. Confocal microscopy and 3D reconstructions of astrocytes revealed that NIF treatment restored astrocyte surface area, volume, and PSD-95 association compared to controls (Figure 7E-H). In summary, these results demonstrate that microglia-mediated pruning of astrocyte processes contributes to the atrophic phenotype observed in astrocytes following LgA cocaine SA and prolonged withdrawal. Furthermore, blocking CR3-C3 mediated phagocytosis during the withdrawal period is sufficient to prevent the onset of the atrophic phenotype and reduce cocaine-seeking behavior.

## DISCUSSION

Prior studies have identified NAc astrocytes as particularly susceptible to drug- and abstinence-induced changes, including impairments in regulation of glutamate homeostasis and decreases in surface area, volume, and synaptic colocalization ^12,13,34^. It has been proposed that a critical aspect of relapse vulnerability following cocaine use is a chronic dysregulation of glutamate homeostasis in the NAc, and this has been attributed to deficits in astrocyte function ^13,35,36^. For example, the glutamate transporter GLT-1 and cystine glutamate exchanger xCT are downregulated in the NAc following cocaine self-administration and abstinence ^37–40^. Moreover, restoring glutamate uptake or exchange by pharmacological intervention decreases cocaine reinstatement ^41–44^. In addition, agonism of astrocyte Gq-coupled DREADD receptors opposes cocaine reinstatement, while knockdown of the astrocyte protein ezrin exacerbates heroin reinstatement ^16,45^. Accordingly, a model has emerged whereby reduced synaptic colocalization of astrocytes leads to compromised glutamate homeostasis, synaptic regulation, and increased drug seeking ^46,47^.

However, despite evidence that NAc astrocytes are structurally smaller and may oppose drug seeking, the mechanisms by which drug self-administration leads to reductions in morphometric features and synaptic colocalization remain unclear. Here we show that abstinence from cocaine self-administration reduces not only overall astrocyte surface area and volume, but also density, number and branching pattern of astrocyte processes, while having no effect on process length. We thus tested the hypothesis that microglial pruning of astrocytes drives decreased structural features of astrocytes in cocaine-abstinent rats. While no overt changes in microglia structural phenotype were observed across abstinence, we observed a time-dependent increase in labeled astrocyte membranes within phagolysosomes in NAc microglia. Further, blockade of complement-dependent microglial phagocytosis during abstinence was sufficient to protect astrocyte structure and synaptic colocalization, and to reduce cocaine seeking behavior. Together, these results demonstrate that microglia pruning drives astrocytic morphological alterations across cocaine abstinence and contributes to cocaine craving.

### Abstinence from cocaine self-administration induces deficits in astrocyte structure

Astrocytes are notably characterized by their complex branching patterns and fine distal processes which comprise approximately 90% of cellular volume and surface area ^48^. A single astrocyte processes can ensheathe a territory of tens of thousands of synapses ^2,5,49^ and enables an astrocyte to detect and modulate synaptic activity ^50,51^. Multiple independent studies have revealed that rat self-administration of cocaine, methamphetamine, and heroin results in decreased synaptic colocalization with astrocytes in the NAc core across extinction or abstinence ^15–17,19^. We found that decrements in astrocyte structure were largest following long-access (LgA, 6 h/day) self-administration and prolonged home cage abstinence, as compared to short-access (ShA, 2 h/day) self-administration and extinction training. While these protocols differ in both degree of cocaine history as well as time and mechanisms of abstinence (home cage versus extinction), they represent two widely employed models of cocaine use and relapse. By reconstructing the 3D branching patterns of individual astrocytes, we found that deficits in astrocyte structure are most apparent in mid to distal regions of the cell (i.e. >20 μm away from the soma) following LgA self-administration and abstinence. Interestingly, we observed decreases in structural complexity independent of our branching classification and that astrocyte processes were on average shorter with fewer branch segments, yet the average distance between any two branch segments was unchanged by cocaine self-administration. We interpret these findings to indicate that reduced morphometric features may not reflect global shrinkage of overall cell size, but rather suggest that a core feature of atrophic astrocytes is a loss of branch processes.

### Microglia pruning of astrocytes during abstinence underlies astrocyte structural alterations and cocaine seeking behavior

Given the known role for microglia in pruning and phagocytosis, we hypothesized that the reduced NAc astrocyte branch complexity could arise as a consequence of aberrant microglia phagocytosis driving astrocyte atrophy. Prior studies indicate that cocaine can directly alter microglia function by activating microglial toll-like receptors (TLRs) ^52–57^ We found that while LgA cocaine self-administration induced a small but persistent increase in the number of microglia within the NAc core, there was no evidence for overt microglial activation, as determined by traditional morphological assessment at either withdrawal day 1 or 45 ^58^. This is in contrast to other studies, which have reported morphological changes early after self-administration ^22,23^. Despite this, we found a robust increase in microglial phagocytosis as evidenced by CD68+ labeling on withdrawal day 45, consistent with the idea that microglia phagocytic function is decoupled from morphological classification ^59^. Furthermore, microglial phagocytosis was associated with engulfment of astrocyte-expressed GFP, which was confirmed using electron microscopy. Together, these data demonstrate that abstinence from cocaine self-administration drives microglial pruning of astrocytes.

It is well-established that microglia prune synapses and circuits in a complement-dependent manner, both in development and pathology ^60–64^. Our data show that complement acts similarly on astrocytes as an “eat me” signal ^65^ during abstinence from cocaine self-administration. Inhibition of the microglial complement receptor 3 during abstinence using Neutrophil Inhibitory Factor (NIF) peptide prevented the decrease in astrocyte structure, restored synaptic colocalization, and decreased drug seeking behavior. The NIF peptide prevents the ability of microglia to “recognize” complement proteins ^64,66^. Interestingly, NIF administration during abstinence did not abolish drug seeking altogether; rather, inhibiting phagocytosis greatly reduced the degree of drug seeking. This finding suggests that microglia-dependent alterations in astrocyte structure may be a key contributor to the incubation of craving that occurs with prolonged abstinence.

Our findings suggest that microglia-induced astrocyte alterations are likely a progressive phenomenon, requiring significant time following cocaine self-administration to fully manifest. This is consistent with other reports that find no changes in astrocyte structure and synaptic colocalization within 24 hours of the last self-administration session ^19^. Similarly, we show here that microglial engulfment of astrocyte material was not increased at 24 hours but was greatly exacerbated at 45 days post self-administration, consistent with the timing of astrocyte alterations ^15,19^. The disparity between the time of cocaine experience and glial pathology poses an interesting question: what is the initiating event that drives microglial engulfment of astrocytes throughout abstinence? We propose two plausible hypotheses: first, that astrocytes may promote their own pruning possibly as an excitotoxic response to the excessive neurotransmitter signaling during cocaine self-administration. To this point, NAc astrocytes are directly responsive to dopamine ^67^ and excess dopamine during self-administration can indirectly increase glutamatergic signaling ^68,69^. In this case, microglia may be responding to cues from astrocytes which results in the eventual pruning of astrocyte processes throughout abstinence. Alternatively, microglia may initiate the pathology with phagocytosis being induced directly by cocaine or by yet unidentified alterations in their microenvironment that result from cocaine self-administration. Further research is needed to better understand exactly how abstinence from cocaine self-administration induces aberrant microglial pruning of astrocytes.

## Conclusion

In conclusion, we demonstrate a novel mechanism contributing to the astrocyte structural deficits that arise from extinction or abstinence following cocaine self-administration and highlight a novel function for microglia in sculpting pathology-induced alterations in brain and behavior. By uncovering new ways in which the neuroimmune system interacts with drugs of abuse, we hope to provide new insight into the biological basis for addiction and relapse. Because deficits in astrocyte structure and function are seen in models of addiction to other drugs of abuse, such as methamphetamine and heroin ^16,17^, it is possible that microglial astrocyte pruning represent a conserved mechanism of pathology within the NAc across drug types.

## Acknowledgements

This work was supported by NIH DA057776 (KJR) and the UNC Cowan Research Excellence Award (KJR).

## Author Contributions

Conceptualization, A.T. and K.J.R.;

Methodology, A.T., J.V.R., T.J.B., and K.J.R.;

Software, M.J.G, H.W..;

Formal Analysis, A.T., J.V.R., and T.J.B.;

Investigation, A.T., J.V.R., T.J.B., R.K., H.A.V., E.A.W., J.P.F, ;

Writing – Original Draft, A.T., J.V.R., and K.J.R.;

Writing – Review & Editing, A.T., J.V.R., T.J.B., and K.J.R.;

Supervision, M-E. T., and K.J.R.;

Funding Acquisition, M-E. T., and K.J.R.

## STAR METHODS

### RESOURCE AVAILABILITY

#### Lead contact

Further information and requests for resources and reagents should be directed to and will be fulfilled by the lead contact, Dr. Kathryn J Reissner.

#### Materials availability

This study did not generate new unique reagents.

#### Data and code availability

All data reported in this paper will be shared by the lead contact upon request. This paper does not report original code. Any additional information required to reanalyze the data reported in this paper is available from the lead contact upon request.

### EXPERIMENTAL MODEL AND STUDY PARTICIPANT DETAILS

#### Animals

Male Sprague Dawley rats were purchased from Envigo (Indianapolis, IN, United States), weighing between 200-250g, and aged 6-8 weeks upon arrival. Rats were individually housed in standard plexiglass cages under controlled temperature and humidity conditions on a reverse light-dark cycle (12:12 h). Rats were allowed to acclimate for a period of one-week, during which rats were allowed ad libitum access to food and water. Subsequently, all rats underwent food restriction (∼20 g chow per day) to facilitate food training and self-administration learning. Food restriction persisted throughout surgical, post-operative, and food-training procedures. Rats were given ad-libitum food during self-administration for the duration of the study once stable lever pressing was achieved. All animal care and procedures were conducted in accordance with the National Institutes of Health *Guide for the Care and Use of Laboratory Animals* and were approved by the Institutional Animal Care and Use Committee at the University of North Carolina.

### METHOD DETAILS

#### Surgical procedures

Animals were anesthetized with ketamine (100 mg/kg) and xylazine (7 mg/kg). A silastic catheter was surgically implanted into the right jugular vein and exited the back attached to a 22G canula (Plastics One), following established protocols ^15,19^. Catheters were flushed daily with antibiotic (gentamicin 5 mg/ml, 0.1 ml i.v.) and heparinized saline (100 U/ml, 0.1 ml i.v.) during post-operative and self-administration procedures. Immediately following jugular vein catheterization, rats were head-fixed in a stereotaxic apparatus (Kopf Instruments) and received microinjections of AAV5-GfaABC1D-Lck-GFP (6.1 x 10^12 virus molecules/ml; Addgene plasmid # 105598; packaged by the University of North Carolina Viral Vector Core) to allow for astrocyte visualization ^70^. Virus was infused bilaterally into the nucleus accumbens (NAc) core (1 μl per hemisphere;6° angle, AP +1.5, ML +2.6, DV -7.2 mm from bregma) at a rate of 0.1 μl/min through 26-gauge guide cannulas (Plastics One, Roanoke, VA). Microinjectors were left in place for 15 minutes post-microinjection to facilitate virus diffusion before a gradual removal over 1-2 minutes. To verify catheter patency before self-administration procedures, a subthreshold dose of propofol (10 mg/mL, 0.05 mL) was administered prior to commencement of self-administration.

For NIF administration experiments, animals also received guide cannula (26G) implants targeting above the NAc core (0° angle, AP +1.4, ML +1.7, DV -5.5 mm from bregma) immediately following virus diffusion. Cannulas were secured in place with the dental cement and wire inserts were placed inside guide cannulas to prevent the entry by contaminants.

#### Self-administration training and behavior

All intravenous self-administration (IV-SA) procedures were conducted in sound-attenuated operant conditioning chambers (Med Associates, St. Albans, VT). Prior to the start of IV-SA, animals underwent a single food-training session, during which responding on the active lever led to the delivery of a single 45 mg food pellet (Bio-Serv). Food training sessions lasted at least 6 hours, with a criterion for learning of over 100 responses on the active lever. Subsequently, rats underwent 10 consecutive days of self-administration, receiving either saline (0.9% NaCl) or cocaine (0.75 mg/kg/infusion) on an FR1 schedule for 2 hours/day (ShA) or 6 hours/day (LgA).Active lever responses triggered the delivery of saline or cocaine (0.045 ml/infusion for a 300 g rat, administered over 2.18 seconds) and were accompanied by a tone (70 dB, 2.5 kHz) and illumination of a stimulus light above the active lever for five seconds. A 20-second timeout followed each infusion, during which active lever responses had no programmed effects. Inactive lever presses at any time during the session had no programmed responses. After the 10-day self-administration period, animals that underwent LgA-SA remained in their home cage for the duration of forced abstinence, while animals that underwent ShA-SA began 15 days of extinction training (2h/day). Extinction training took place in the same operant chambers. During extinction training sessions, levers were extended but presses did not elicit drug infusions or sound/light cues.

For NIF administration experiments, all rats underwent LgA cocaine self-administration. Animals received bilateral microinfusions of Neutrophil Inhibitory Factor (NIF) peptide (200 μg/mL; 0.5 μL/hemisphere at 0.1 μL/min; 1 min diffusion time) or vehicle (sterile PBS), beginning 24 hours after the last LgA-SA session and repeated every 7 days for a total of 4 NIF peptide infusions throughout abstinence. Forty-eight hours after the fourth infusion (Withdrawal Day 24; WD24), animals underwent a drug-seeking test in the same operant chambers where active lever presses elicited drug-associated cues (e.g. light, tone) but did not result in drug infusion.

#### Tissue collection and immunohistochemical processing

Animals were euthanized either 24 hours (WD1), 45 days (WD45) or 24 days (NIF experiment; WD24) after the last LgA-SA session. Animals undergoing ShA-SA and extinction were sacrificed 24h after the last extinction training day. Animals were anesthetized with sodium pentobarbital, followed by transcardial perfusion with 1x phosphate buffer (PB) and 4% paraformaldehyde (PFA) in PB. Brains were extracted, post-fixed in 4% PFA for ∼4 hours, and stored in 30% sucrose. Coronal sections (45 and 100 μm) from the NAc were obtained using a cryostat (Leica) and stored in 50% glycerol/PBS until immunohistochemical processing.

For immunohistochemistry of thick sections (100 μm), free-floating NAc sections were washed (3 x 5 min) in 1x PBS containing 2% Triton X-100 (PBST-2%) (Thermo Fisher, Waltham, MA). Subsequently, sections were blocked in 5% normal goat serum (NGS, Sigma Aldrich, St. Louis, MO) in PBST for 1 hour at room temperature. Sections were then incubated with primary antibodies (mouse anti-PSD-95 (Thermo Fisher, Waltham, MA) and rabbit anti-GFAP (Dako, Santa Clara, CA), both at 1:500) in 5% NGS-PBST for 72 hours at 4 °C and turned around halfway through the incubation for optimal penetration. Following primary antibody incubation, sections were washed (3 x 5 min) with PBST, then incubated with secondary antibodies (goat anti-mouse Alexa Fluor 594 (Thermo Fisher, Waltham, MA) and goat anti-rabbit Alexa Fluor 647 (Thermo Fisher, Waltham, MA), both at 1:1000 in 5% NGS-PBST) for 72 hours at 4 °C with a tissue being turned around half-way through. After secondary antibody incubation, sections were washed (3 x 10 min in 1x PBS), mounted onto slides, and coverslipped with DAPI Fluoromount-G (Southern Biotech, Birmingham, AL).

For immunohistochemistry of thin sections (45 μm), free-floating NAc sections were washed (3 x 5 min) in 1x PBS containing 0.2% Triton X-100 (PBST-0.2%) followed by blocking with 5% NGS-PBST for 1 hour at room temperature. Sections were incubated with primary antibodies (mouse anti-CD68 (1:400), rabbit anti-Iba1 (1:1000), chicken anti-C3 (1:500) in 5% NGS-PBST-0.2% overnight at 4 °C. The next day, sections were washed(3 x 5 min) in PBST-0.2, then incubated with secondary antibodies (goat anti-mouse Alexa Fluor-633, goat anti-rabbit Alexa Fluor-405, goat anti-chicken Alexa Fluor-594 in 5% NGS-PBST-0.2%) for 2 hours at room temperature. Sections were then washed (3 x 10 min) in PBS, mounted onto slides and coverslipped using DAPI-free Fluoromount-(Southern Biotech, Birmingham, AL).

#### Confocal image acquisition

Image acquisition and processing of NAc astrocytes using thick sections (100 μm) followed previously described methods (Testen et al., 2019; Testen et al., 2018, also see Testen et al. 2020 for detailed protocol). Briefly, imaging was performed using a Zeiss LSM 800 confocal-scanning microscope equipped with 405, 488, 561, and 640 lasers. Images were acquired using 63x (1.4 NA) oil-immersed objective with a 1024 x 1024 pixels field of view (FOV) at 16-bit depth, 4x averaging, and 1 μm z-step. Only single, isolated astrocytes within the NAc core were imaged, excluding those that bordered other astrocytes or were cut within the z-plane during sectioning. Microglia images were acquired from thin sections using identical parameters and a 0.5 μm z-step to increase resolution along the z-axis. Microglia were randomly sampled within the NAc core regardless of their proximity to astrocytes. Following acquisition, raw images of both cell types were deconvolved (blind; 10 iterations) using AutoQuant software (v. X3.0.4, MediaCybernetics) with resulting images being saved as Imaris files (.ims).

#### Astrocyte surface reconstruction, synaptic colocalization, and complexity analysis

Astrocyte 3-dimensional surface reconstructions were created using Imaris software (v 8.4.1 and 10.1, Bitplane, Zurich, Switzerland). Using the Lck-GFP signal from each astrocyte, the Surfaces module was used to create a volumetric boundary of each cell. The resulting surface was used to calculate surface area and volume for each astrocyte and was used as a mask to determine PSD-95 colocalization, which was used as a proxy for post-synaptic terminals. The background threshold for the PSD-95 signal was manually determined by averaging several measurements of clear PSD-95 signals across each z-stack. A colocalization channel was generated to determine the percentage of PSD-95 signal colocalized within the masked Lck-GFP astrocyte signal.

Individual astrocytes were reconstructed with the Filaments module using the “Autopath” feature which employs the Fast-marching method (Sethian, 1996), using the astrocyte mask channel (described above) as a reference. The maximal number of filament seed points was determined by the lateral resolution of the image (297 nm), and DAPI-stained nuclei of each individual astrocyte served as the starting point for the filament construction, which was verified by colocalization with both GFAP labeling and Lck-GFP signal for each astrocyte. Reconstructed astrocyte filaments served as the input for all further complexity analyses within Imaris software, namely 3D Sholl analysis, branching analysis, analysis of filament terminal points, and segment analysis. The 3D Sholl analysis (XTension: Filament Sholl Analysis; ImarisOpen depository) diameter was set at 1 µm and included the number of Scholl intersections, the peak intersection number, distance of the peak intersection number, and area under the curve (AUC) of the plotted distribution. The branching analysis (XTension: Filament to Dendrogram; ImarisOpen depository) was used to identify and classify filament bifurcation nodes, filament endings (terminal points), and segment length. Bifurcation nodes were classified as either arborizing (both child branches continue to branch past the node), continuing (one child branch continue to branch while the other terminates), or terminating (both child branches terminate past the node) and their distribution was analyzed similarly to the 3D Sholl analysis. Filament terminal points were used as a proxy for peripheral astrocyte processes (PAPs). A segment was defined as a part of the astrocyte filament between two distinct bifurcation nodes.

#### Microglia inclusion and complexity analysis

Microglia 3-dimensional surface reconstructions were performed similarly to astrocytes using Imaris software, by isolating the Iba1 channel mask for each individual microglia in the z-stack. The Surfaces module was used to create a surface for each microglia and calculate surface area and volume. The Iba1 channel mask was used to determine colocalization for each of the C3, CD68, and Lck-GFP signals. Signal threshold for each channel was determined as described above for PSD-95/astrocyte colocalization, with fluorescence intensity measurements performed for multiple distinct, unambiguous puncta of each channel. The average intensity value was then used as a threshold and the resulting colocalization was reported as percentage of each microglia cell volume.

To analyze potential astrocytic inclusions within microglia, the Iba1/Lck-GFP colocalization channel was used to build a 3-dimensional surface and the resulting output was filtered to eliminate all surface objects smaller than the limit of resolution of all dimensional axes (0.25 μm x 0.25 μm x 0.5 μm = 0.03125 μm^3^). The volumes of all surface objects were then summed to obtain the total volume of inclusions per cell, and the volume of soma inclusions was calculated by only summing surfaces within the reconstructed soma. Quantification of Lck-GFP-positive signal within microglial phagosomes (CD68+) was performed by colocalization analysis between the Iba1/Lck-GFP mask (i.e. total astrocyte inclusions) and the Iba1/CD68 reference mask (i.e. total phagosomes). Inversely, the quantification of the percentage of inclusions contained within phagosomes was performed by colocalization analysis between the Iba1/CD68 mask and the Iba1/Lck-GFP reference mask.

Individual microglia were reconstructed with the Filaments module using the Iba1 mask channel (described above) as a reference. Each microglia soma was selected as a starting point for the filament using the “automatic soma filling” feature whenever possible to standardize reconstructions. In the few cases where automated soma filling failed to produce an appropriate soma, each soma surface was built manually in conjunction with the ROI module. The resulting soma surface was then imported during the filament building process as a starting point and was used to quantify the area, volume and soma sphericity Each microglia filament reconstruction was then analyzed using the Sholl Intersections module to quantify microglia complexity, and the Convex Hull XTension was used to determine the volume of “functional” territory occupied by each microglia.

#### Quantification of microglia number

Coronal sections (45 µm thick) were immunolabeled for Iba1 and imaged via confocal microscopy as described above. Images were acquired within the NAc core using a 63x objective with a 2×2 tile scan and 1 μm z-steps through the entire tissue thickness, and the resulting z stack was used for cell counting using Imaris software. At least two fields of view were imaged across two sections for each animal, and the number of Iba1+ cells was quantified. Cells were included in the analysis if a complete soma could be identified for each cell.

#### Tissue collection for scanning electron microscopy

Brain sections were prepared for scanning electron microscopy as previously described ^71^. Briefly, animals were euthanized 45 days after the last LgA-SA session (WD45). Animals were anesthetized with sodium pentobarbital, followed by transcardial perfusion with 3.5% acrolein in PB, then with 4% PFA in PB. Brains were extracted and post-fixed in 4% PFA for 2 hours. After post-fixation, brains were washed 3 x 10 min in PBS and coronal sections of the NAc (50 μm thick) were prepared using a vibratome. Sections were stored in cryoprotectant at -20°C until processed.

#### Immunostaining, processing, and scanning electron microscopy

Brain sections containing the NAc core were processed for immunostaining against Lck-GFP prior to scanning electron microscopy (SEM) processing. Briefly, samples were washed with phosphate buffered saline (PBS; pH 7.4) (5 x 7 min at RT), exposed to citrate buffer for antigen retrieval (15 min at 70 °C), washed with PBS (3 x 7 min at RT), quenched with 0.3% H_2_O_2_ in PBS (5 min at RT), washed with PBS (3 x 7 min at RT), incubated in 0.1% NaBH_4_ in PBS (30 min at RT), washed with PBS (3 x 7 min at RT) before incubation in a blocking buffer solution (5 % normal goat serum, 3 % bovine serum albumin, 0.1 % Triton-X-100 in tris buffered saline (TBS; 50 mM, pH 7.4) (1 h at RT). Sections were incubated overnight at 4 °C in blocking buffer solutions with rabbit anti-GFP IgG antibody (1:500) (Invitrogen). The next day, sections were washed with TBS (3 x 7 min at RT), incubated with a biotinylated goat anti-rabbit polyclonal antibody (1:300) (JacksonImmunoResearch) in blocking buffer (90 min at RT), washed with TBS (3 x 7 min at RT) and incubated with an avidin-biotin complex solution (1:100) in TBS (1 h at RT) and washed with TBS (3 x 7 min at RT). The staining was revealed with 0.05% 3,3′-diaminobenzidine (activated with 0.015% H_2_O_2_ diluted in Tris buffer (0.05 M, pH 8.0) for 1 min before being stopped by transferring to phosphate buffer (PB) for washes (3 x 7 min) and stored overnight at 4 °C.

The following day sections were rinsed in PB (3 x 5 min) and incubated in a solution containing equal volumes of 3% potassium ferrocyanide and 4% osmium tetroxide in PB (1 hr at RT). Sections were then washed in PB (1 x 5 min at RT), 1:1 PB and MilliQ water solution (1 x 5 min at RT) and MilliQ water (1 x 5 min at RT) before incubation in 1% thiocarbohydrazide in MilliQ water (20 min at RT). Sections were then washed in MilliQ water (3 x 5min at RT) before incubation with 2% aqueous osmium tetroxide in MilliQ water (30 min at RT), after which the sections were incubated in MilliQ water (5 x 5 min at RT). The sections were dehydrated in ascending concentrations of ethanol, 2 × 35%, 50%, 70%, 80%, 90%, 3 × 100% (5 min each at RT), and then washed in propylene oxide (3 x 5 min at RT) prior to embedding in Durcupan resin (10 g component A, 10 g component B, 0.3 g component C, 0.2 g component D) overnight (at RT). The next day, the resin-infiltrated sections were flat-embedded between two fluoropolymer ACLAR® films and polymerized (3 days at 55 °C).

From the polymerized sections, areas containing the nucleus accumbens core were excised and glued onto resin blocks for ultra-thin sectioning. Using a Leica ARTOS 3D ultramicrotome, 70-nm thick sections were made at multiple levels (∼5–10 μm apart) and collected on silicon wafers. Ultra-thin sections were imaged on a Zeiss Crossbeam 350 focused ion beam-scanning electron microscope (FIB-SEM). Using Zeiss Atlas 5 software, images of microglia were acquired in the NAc core at a resolution of 5 nm per pixel and exported in .tif format.

### QUANTIFICATION AND STATISTICAL ANALYSIS

Statistical analysis was performed using GraphPad Prism (version 10). Data are presented as the mean ± SEM along with individual data points. Additional statistical details (tests used and statistic, exact n, p values, etc) can be found in figure legends and text. Group sizes were not predetermined statistically but are consistent with those previously reported in the field. A p value of < 0.05 was used as the criterion for significance. All behavior, imaging, and analysis was performed by experimenters blind to treatment and condition.

Two- or three-way repeated measures analysis of variance (ANOVA) was used when appropriate to analyze self-administration and extinction training data, astrocyte complexity, and microglia complexity data. Statistically significant interactions were further analyzed using Bonferroni’s post-hoc tests to determine exact differences. NIF administration data was not normally distributed based on the D’Agostino & Pearson distribution test and was therefore analyzed using Mann-Whitney test for significance. For comparisons of astrocyte morphological parameters between 4 treatment groups independent of distance, a one-way nested ANOVA was used with Tukey’s multiple comparisons post-hoc test using animal as a main factor and individual cell values as a nested factor. For comparisons of microglial measurements between two groups independent of distance, a nested two-tailed t-test was used. Area under the curve (AUC) was compared using unpaired two-tail t-test.

Principle components analysis (PCA) was performed using morphological (astrocytes and microglia) or inclusion-related variables (microglia). Variables were averaged per animal and used as independent continuous variables. Results and graphs for loadings, eigenvalues, and proportions of variance can be found in supplemental material. Principial component regression (PCR) analysis using least squares regression type was conducted afterward where principal components of each morphological or inclusion-related variable were correlated to the corresponding behavioral result for each animal (number of infusions or active lever presses). Principal components used in PCR were selected using the Kaiser rule (eigenvalue >1.0). Regression correlations deemed statistically significant were confirmed by a simple linear regression using raw averages (instead of principal components used in PCR).

